# Discovery of a new species of subterranean eel loach from southern India

**DOI:** 10.1101/2024.09.10.612189

**Authors:** K.R. Sreenath, B. Pradeep, K. R. Aju, Sandhya Sukumaran, Wilson Sebastian, Alvin Anto

## Abstract

*Pangio juhuae* sp nov, a new species of subterranean eel loach, is described from Kerala, India. It is distinguished from its subterranean congeners by the presence of a dorsal fin, four pectoral rays and five segmented anal fin rays. Genetic analyses suggest that *P. juhuae* is closely related to *P. bhujia* but is distinct in morphology, particularly by the presence of a dorsal fin. The low genetic distance and significant morphological difference between these two *Pangio* species suggest that they have diverged from an immediate common ancestor and evolved distinct adaptations to subterranean niches. *P. juhuae* exhibits less evolved troglobitic traits compared to *P. bhujia* and *P. pathala*, suggesting it could be a connecting species in the evolutionary transition from terrestrial to subterranean loaches. This discovery provides evidence for possible subterranean speciation of fishes in underground habitats.

## Introduction

Troglobitic or stygobitic fishes, found in subterranean habitats worldwide except Antarctica, exhibit troglomorphism, adaptations to life in caves and underground cavities. Recent years have seen a surge in the discovery of new subterranean fish species, with the global count reaching 309 cave and groundwater species. Kerala, in southwestern India, emerges as a potential hotspot for subterranean fish diversity, boasting 12 described species from four families across four orders. The Cobitidae family, specifically the genus Pangio (eel-loaches), is less common in underground habitats globally compared to Cyprinidae. The present study focuses on five specimens of a subterranean eel-loach collected from a dugout well in Naduvannur, Kozhikode, northern Kerala, potentially representing a new species.

Four specimens of *Pangio juhuae* sp. nov. were collected and preserved for morpho-meristic and molecular analysis. Morphometric data were recorded, and vertebrae and fin rays were counted. Another specimen was used for molecular study. DNA was extracted using a standard phenol/chloroform protocol, and a 650 bp region of the Cytochrome C oxidase 1 gene was amplified and sequenced. The COI sequence was deposited in GenBank. Phylogenetic analysis was conducted. Genetic divergence between *Pangio juhuae* and other Pangio species was analyzed using Kimura 2 p distance values and uncorrected p values.

## Results

### Descriptions of the new species

Class: *Teleostei*

Order: *Cypriniformes*

Family: *Cobitidae* Swainson, 1838

Genus: *Pangio* Blyth, 1860

Species: *Pangio juhuae* sp. nov.

Type Material: *Holotype:* CMFRI DNR GB 22.4.1.1; *Paratypes:* (I) CMFRI DNR GB 22.4.1.1.1, (II) CMFRI DNR GB 22.4.1.1.2

Suggested Common Name: Juhu’s pigmy eel-loach

### Diagnosis

*Pangio juhuae* is a distinct species among its hypogeal congeners due to the presence of a prominent dorsal fin. It has a relatively larger eye diameter compared to *P. bhujia* and *P. pathala*. It differs from *P. bhujia* in having four pectoral fin rays (vs. three) and five anal fin rays (vs. six). Trogomorphic traits such as a slender body, lack of pigmentation, scales, and diminutive eyes differentiate it from other non-subterranean congeners.

### Description

The fish species described in the text has a thin, elongated, and anguilliform body with a small, rounded head. It lacks scales and possesses four pairs of barbels. The dorsal fin is located in the posterior half of the body and has five rays. The pectoral fin is slender with four segmented rays, while the pelvic fin is absent. The anal fin is short and rounded with five unbranched and segmented rays. The caudal fin is lanceolate with 14 principal rays and 6 procurrent rays, all segmented. The vertebrae count, estimated using microCT, was Webberian + 57 + Hypurals. It has a translucent body, vertebrae visible, abdominal region burnt red due to internal organs and blood vessels, reddish brown on the dorsal side, transparent tan caudal peduncle, caudal artery visible as a red streak, diminutive black specks for eyes. Beige in preservative, dark spots for eyes. This information provides a comprehensive overview of the fish’s morphological characteristics. The species is currently known only from the type locality, Naduvannur, Kerala, India. It was first observed in a water tank filled with water from a dug-out well, supplied by underwater springs. It is named after a four-year-old girl, Juhu, who first noticed it.

**Table 1.**
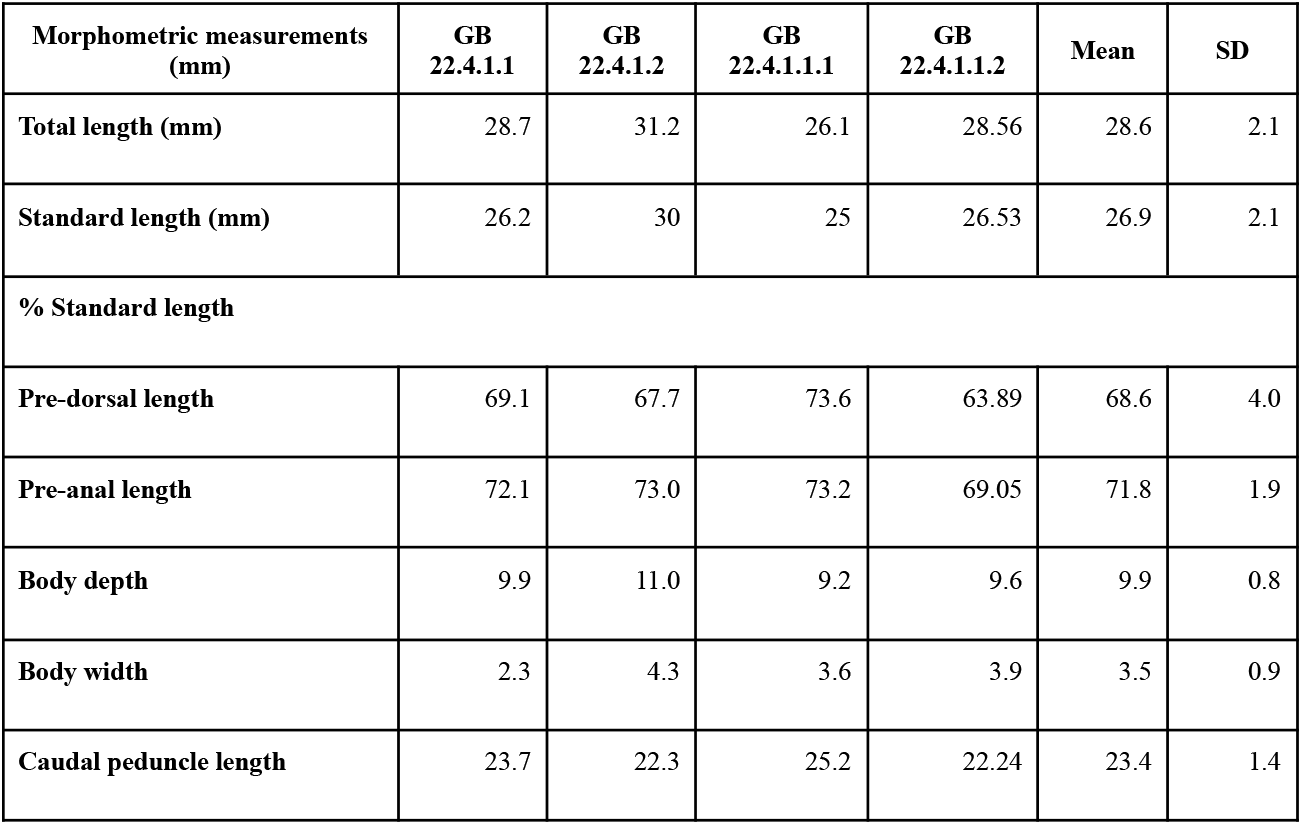
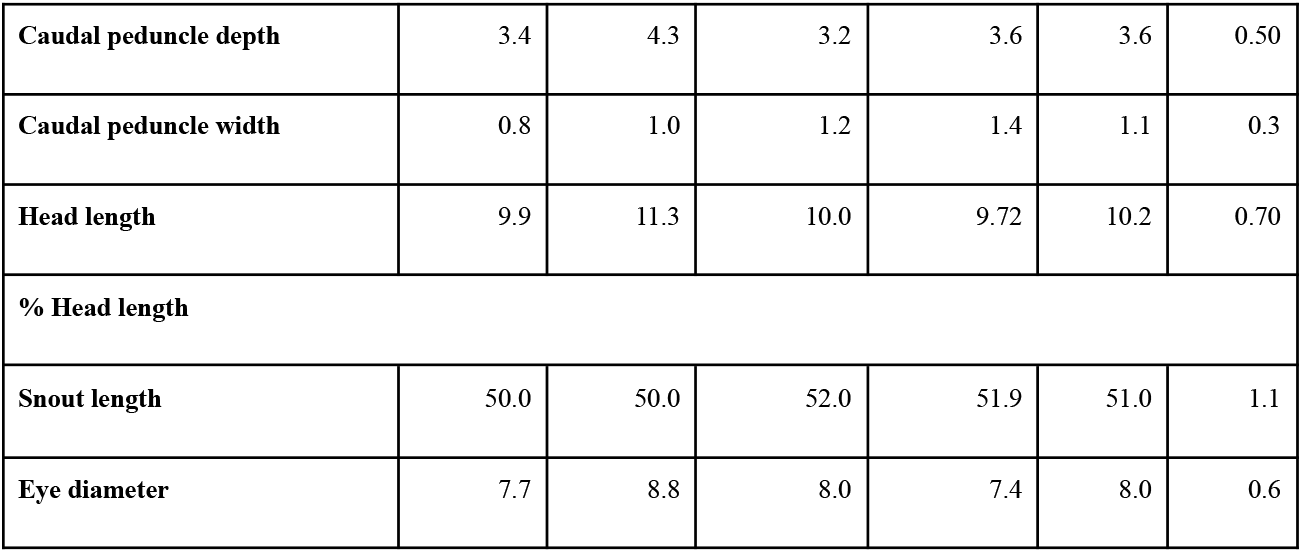
Morphometric measurements of *Pangio juhuae*.

| Morphometric measurements (mm) | GB 22.4.1.1 | GB 22.4.1.2 | GB 22.4.1.1.1 | GB 22.4.1.1.2 | Mean | SD |
| --- | --- | --- | --- | --- | --- | --- |
| Total length (mm) | 28.7 | 31.2 | 26.1 | 28.56 | 28.6 | 2.1 |
| Standard length (mm) | 26.2 | 30 | 25 | 26.53 | 26.9 | 2.1 |
| % Standard length |  |  |  |  |  |  |
| Pre-dorsal length | 69.1 | 67.7 | 73.6 | 63.89 | 68.6 | 4.0 |
| Pre-anal length | 72.1 | 73.0 | 73.2 | 69.05 | 71.8 | 1.9 |
| Body depth | 9.9 | 11.0 | 9.2 | 9.6 | 9.9 | 0.8 |
| Body width | 2.3 | 4.3 | 3.6 | 3.9 | 3.5 | 0.9 |
| Caudal peduncle length | 23.7 | 22.3 | 25.2 | 22.24 | 23.4 | 1.4 |
| **Caudal peduncle depth** | 3.4 | 4.3 | 3.2 | 3.6 | 3.6 | 0.50 |
| **Caudal peduncle width** | 0.8 | 1.0 | 1.2 | 1.4 | 1.1 | 0.3 |
| **Head length** | 9.9 | 11.3 | 10.0 | 9.72 | 10.2 | 0.70 |
| **% Head length** |  |  |  |  |  |  |
| **Snout length** | 50.0 | 50.0 | 52.0 | 51.9 | 51.0 | 1.1 |
| **Eye diameter** | 7.7 | 8.8 | 8.0 | 7.4 | 8.0 | 0.6 |

**Fig. 1.**
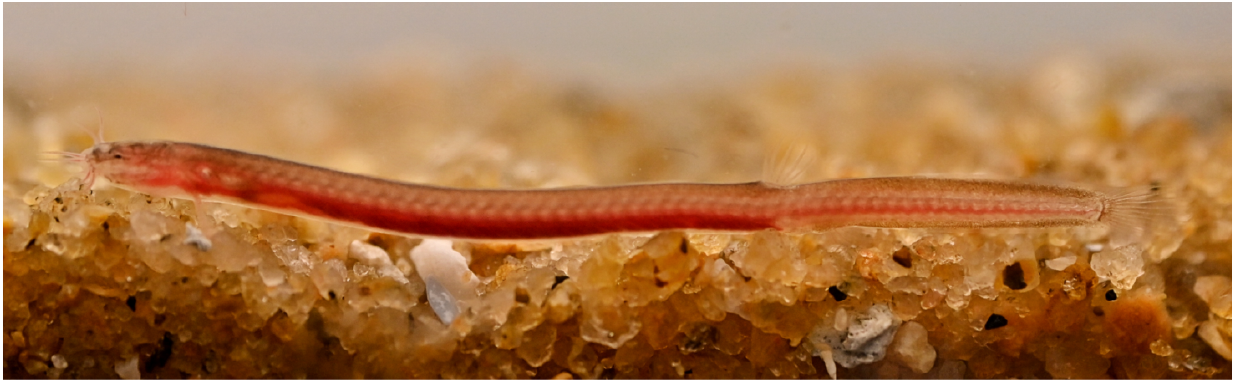
*Pangio juhuae*; Holotype: CMFRI DNR GB 22.4.1.1; Type Locality: Naduvannur, Kozhikode District, Kerala, India.

**Fig 2.**
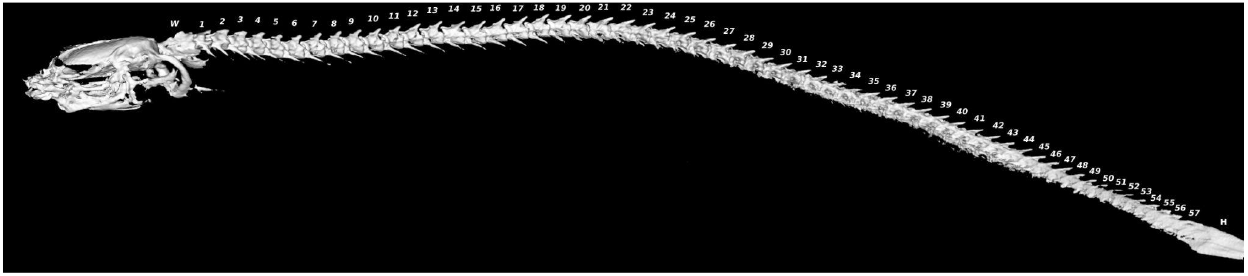
MicroCT image of *P. juhuae*. W: Weberian apparatus, Numbered Vertebrae (57), H: Hypural.

## Discussion

The Cobitidae family, particularly the genus Pangio, presents a compelling subject for biogeographic studies. The discovery of subterranean species within *Pangio* in Kerala adds to the complexity of underground habitats and biodiversity. Three hypogean *Pangio* species in Kerala (*P. bhujia, P. pathala*, and *P. juhuae*) exhibit regressive troglomorphic traits adapted to their underground habitats. *Pangio juhuae* stands out for its intermediate troglomorphic features, suggesting it might still be evolving towards a fully hypogean life. Phylogenetic distance between *P. juhuae* and *P. bhujia* exceeds the lowest inter-species variation among congeners, indicating recent divergence and adaptation to distinct subterranean niches. The presence of a dorsal fin could be a remnant of its surface-dwelling ancestry, gradually disappearing as it adapts to life underground.

Furthermore, this finding potentially exemplifies subterranean speciation. The unique challenges of subterranean environments, such as darkness, limited food, and specific water flow, exert distinct selective pressures that can lead to rapid morphological and genetic changes, potentially driving the formation of new species even within a short timeframe. The discovery of this loach, with its unique dorsal fin and close genetic relationship to other subterranean species, illustrates how adaptation to these niche environments can facilitate speciation. Further research into its morphology, genetics, and behavior could unveil the adaptive significance of the dorsal fin in its subterranean habitat and identify the specific genetic changes associated with the transition to subterranean life. Comparing its genome to other Pangio species, especially *P. bhujia*, may pinpoint genes involved in fin development, sensory perception, or metabolism that play a role in this adaptation process. Additionally, behavioral observations could reveal further adaptations to the underground environment.

In conclusion, this discovery could serve as a valuable model for understanding subterranean speciation and the adaptive mechanisms underlying the colonization of these unique habitats. Further research into this loach has the potential to shed light on broader evolutionary patterns and the remarkable biodiversity thriving in the hidden world beneath our feet.

## Acknowledgements

The authors acknowledge the Director, ICAR-Central Marine Fisheries Research Institute & the Director, ICAR-Indian Institute of Spices Research for the support. The authors also thank Ms Aswani, Naduvannur for their help with the collection of the specimens and Mr Rahul G. Kumar for improving the manuscript with valuable suggestions. We also wish to appreciate the facilities extended by Amrita Institute of Medical Sciences for imaging the specimens.

